# Genomic diversity and antimicrobial resistance in clinical *Klebsiella pneumoniae* isolates from tertiary hospitals in Southern Ghana

**DOI:** 10.1101/2024.01.20.576413

**Authors:** Richael O. Mills, Isaac Dadzie, Thanh Le-Viet, David J. Baker, Humphrey P. K. Addy, Samuel A. Akwetey, Irene E. Donkoh, Elvis Quansah, Prince S. Semanshia, Jennifer Morgan, Abraham Mensah, Nana E. Adade, Emmanuel O. Ampah, Emmanuel Owusu, Philimon Mwintige, Eric O. Amoako, Anton Spadar, Kathryn E. Holt, Ebenezer Foster-Nyarko

## Abstract

Comprehensive data on the genomic epidemiology of hospital-associated *Klebsiella pneumoniae* in Ghana is scarce. This study sequenced 103 clinical *K. pneumoniae* isolates from five tertiary hospitals in Southern Ghana, predominantly from paediatric patients under five years (67/103, 65%), with the majority collected from urine (32/103, 31%) and blood (25/103, 24%) cultures. We employed Pathogenwatch for genotyping via Kaptive (K/O antigens) and Kleborate (antimicrobial resistance and hypervirulence) and determined clonal relationships using core-genome multilocus sequence typing (cgMLST). Among the 44 distinct sequence types (STs) detected, ST133 was the most common, comprising 23% of isolates (n=23/103). We discovered 27 different capsular (K) locus antigens and seven lipopolysaccharide (O) types; KL116 (28/103, 27%) and O1 (66/103, 64%) were the most prevalent. Single-linkage clustering highlighted the global spread of multidrug-resistant clones such as ST15, ST307, ST17, ST11, ST101, and ST48, with minimal allele differences (1-5) from publicly available genomes worldwide. Conversely, several isolates (n=17) constituted novel clonal groups and lacked close relatives among publicly available genomes, displaying unique genetic diversity within our study population. A significant proportion of isolates (88/103, 85%) carried resistance genes for three or more antibiotic classes, with the *bla*_CTXM-15_ gene present in 78% (n=80/103). Carbapenem resistance, predominantly due to *bla*_OXA-181_ and *bla*_NDM-1_ genes, was found in 10% (n=10/103) of the isolates. Yersiniabactin was the predominant acquired virulence trait, identified in 70% (n=72/103) of the isolates. Our findings reveal a complex genomic landscape of *K. pneumoniae* in Southern Ghana, underscoring the critical need for ongoing genomic surveillance to manage the substantial burden of antimicrobial resistance.

## Introduction

*Klebsiella pneumoniae* is notoriously linked with multidrug-resistant infections, particularly in healthcare settings [1–8]. Within sub-Saharan Africa, *K. pneumoniae* has emerged as the second most frequent causative agent and leading Gram-negative agent in neonatal sepsis cases, underscoring a pressing need for effective surveillance and control measures [9]. The Global Research on Antimicrobial Resistance study estimated that drug-resistant *K. pneumoniae* contributed to over 600,000 deaths in the region in 2019 [4]. Thus, *K. pneumoniae* represents a pressing health challenge in the current era of increasing antimicrobial resistance (AMR).

In Ghana, the clinical impact of *K. pneumoniae* is well-documented, with increasing resistance to critical antibiotics like third-generation cephalosporins and carbapenems being a notable concern locally [10–21], as it is globally [22]. However, relatively little is known about the pathogen variants underlying this clinical problem [10, 12–14, 16–21].

Despite the paucity of comprehensive local molecular epidemiological data [15, 16, 21, 23, 24], the information available paints a grim picture of the extensive challenges posed by this pathogen’s persistent and complex AMR mechanisms. For example, one study observed a 41% prevalence of gut colonisation with ESBL-producing *K. pneumoniae* in 435 children under five in the Agogo municipality, pinpointing a community-wide reservoir of the *bl*a_CTX-M-15_ gene [15]. Furthermore, research into poultry meat contamination in Kumasi, Ghana, reported 18% of the poultry meat samples tested to carry ESBL *K. pneumoniae*, with *bla*_CTX-M-15_ as the predominant gene [25]—raising alarm bells about the zoonotic transmission of resistant strains.

A One Health study from Northern Ghana highlighted a 64% prevalence of AMR genes in clinical settings, including the discovery of two carbapenemase-producing isolates (an ST17 clone with *bla*_OXA-181_ and an ST874 carrying a *bla*_OXA-48_), emphasising the need for targeted intervention [21]. Moreover, the genomic complexity observed in the Komfo Anokye Teaching Hospital in Kumasi, Ghana, featuring diverse resistance genes on mobile plasmids, highlights the escalating threat of multidrug resistance in hospital-acquired infections [15, 23]. Notably, plasmids carrying replicons such as IncF, IncX3, and IncL have been implicated in disseminating critical resistance genes, such as *bla*_CTX-M-15_, *bla*_OXA-181_, and *bla*_OXA-48_, without a fitness cost, in *K. pneumoniae* and *K. quasipneumoniae* isolates from Effia Nkwanta Hospital, Ghana, underscoring their potential for widespread transmission [24].

WGS has revolutionised pathogen surveillance by providing intricate details of pathogen characteristics, evolution, and transmission pathways. However, the resolution offered by conventional short-read technologies like Illumina is often insufficient for thoroughly resolving plasmids and mobile genetic elements, pivotal for a comprehensive understanding of AMR dynamics. In contrast, long-read sequencing platforms such as the Oxford Nanopore MinION offer the capability to unravel complex genetic architectures [26–28], with hybrid Illumina-Nanopore assemblies leveraging the strengths of both technologies [29–31].

The deployment of WGS in regions with limited resources, such as Ghana, faces barriers, including cost and technical expertise [32–34]. Initiatives like SEQAFRICA are mitigating these challenges by offering sequencing services to investigate AMR related questions. Still, the necessity for local sequencing and analytical capabilities remains to ensure rapid and relevant responses to AMR [35].

Here, we utilised a collection of *K. pneumoniae* isolates from major referral hospitals in Southern Ghana to examine the population structure and AMR transmission dynamics in a high-risk setting. By integrating Nanopore and Illumina data, we constructed hybrid reference assemblies to elucidate the genomic diversity and antimicrobial resistance profiles of clinical *K. pneumoniae* isolates from tertiary healthcare facilities in Southern Ghana.

## Methods

### Sample population and isolate recovery

From January 2021 through October 2021, we prospectively collected *Klebsiella pneumoniae* isolates identified via routine diagnostics from four regions in Ghana: Greater Accra, Ashanti, Central, and Western (**Figure 1A**). The participating facilities included Korle-Bu Teaching Hospital (KBTH) in Accra, the largest tertiary hospital in Ghana, with a 2000-bed capacity; Greater Accra Regional (Ridge) Hospital, also in Accra, serving as a secondary level facility for the Greater Accra Region; Komfo Anokye Teaching Hospital (KATH) in Kumasi, the second-largest with 1200 beds; Cape Coast Teaching Hospital (CCTH) with 400 beds; and Effia Nkwanta Regional Hospital in Sekondi-Takoradi, a significant secondary facility. Blood culture requests were made for all patients with suspected sepsis, although some patients opted to utilise neighbouring private laboratories. Other samples were derived from routine diagnostic processes. Clinical data accompanying these isolates were retrieved from the laboratory records of these hospitals. Additionally, historical isolates from KATH, dating between October 2017 and May 2018, were included in the analysis.

**Figure 1.**
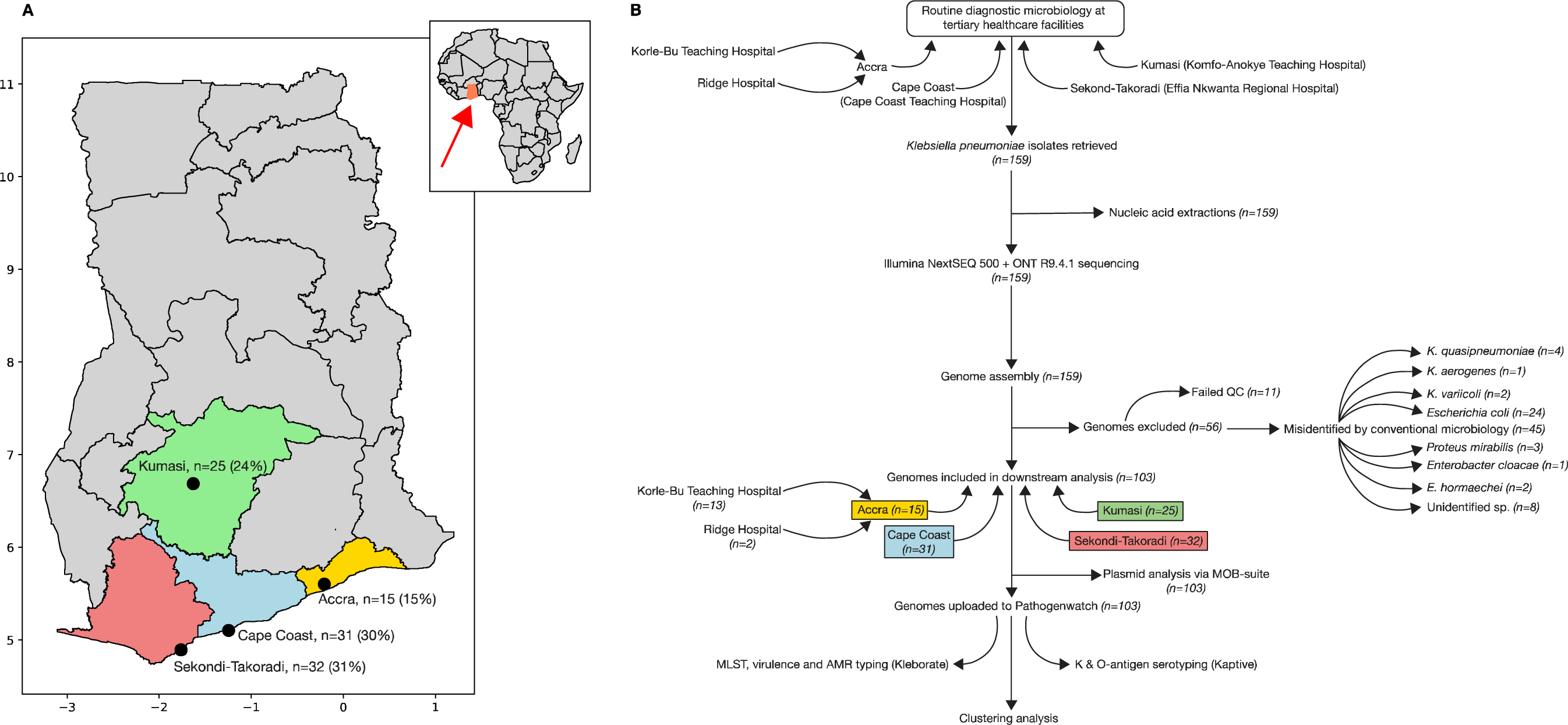
Geographic overview of Study sites in Ghana and the Study sample processing flow. **Panel A:** A map showcasing the Study sampling sites in Ghana, with the different Regions where these cities are located highlighted in gold (Greater Accra), light-green (Ashanti), light-blue (Central) and light-coral (Western). The number of samples derived from each sampling site is shown as a proportion of the total. The inset depicts Africa with Ghana highlighted in coral (red arrow). **Panel B:** The study sample processing flow diagram. The flowchart illustrates the methods utilised for the isolation and genomic characterisation of the study isolates, spanning sample collection, culture and isolation, genomic DNA extraction, whole genome sequencing and analysis aimed at identifying key genetic features, such as antimicrobial resistance markers and virulence factors.

Isolation and identification of *K. pneumoniae* were conducted using the following procedures. Swab, sputum, wound, and aspirate samples were initially cultured on MacConkey agar, Columbia agar supplemented with sheep blood, and chocolate agar plates. These plates were then incubated at 37°C overnight—the chocolate agar in a 5% CO2 atmosphere and the others aerobically. For blood cultures, specimens were inoculated into either brain-heart infusion or Mueller-Hinton broth within a universal or BD BACTEC (Becton, Dickinson and Company, UK) bottle, adhering to a blood-to-broth ratio between 1:10 and 1:20. The cultures were incubated at 37°C for five days or up to 21 days for suspected cases of endocarditis, with daily visual inspections. Blood culture bottles exhibiting turbidity, a sign of bacterial growth, were subjected to subculturing on MacConkey agar, Columbia agar with sheep blood, and chocolate agar, and incubated as previously mentioned. Urine samples were streaked on Cysteine Lactose Electrolyte-Deficient (CLED) agar plates divided into quadrants, using a 10 µl sterile loop for a single streak per quadrant.

Post-incubation, plates were examined for growth characteristics indicative of *Klebsiella* species, as follows. On blood or chocolate agar, colonies with shiny, mucoid, and round appearance with a greyish-white hue and a diameter greater than 3 mm, typical of *Klebsiella* were selected. Strains exhibiting a unique “cotton candy” or “stringy” morphology due to an extensive capsular layer were included. On MacConkey agar, lactose-fermenting colonies with a smooth, mucoid, and pink to reddish appearance were picked. On CLED agar, candidate raised, mucoid colonies with a yellow coloration, denoting lactose fermentation were selected for further processing.

Putative *Klebsiella* isolates were further tested biochemically using indole, citrate, urea, and triple sugar iron (TSI) assays. In brief, a single colony-forming unit (CFU) was inoculated into peptone water for the Indole test, while for citrate, urea, and TSI tests, a CFU was introduced into the respective media via a stab inoculation followed by streaking on the surface slant. These tests were incubated at 37°C overnight. Isolates that were urease and citrate positive, indole negative, and demonstrated glucose fermentation with gas production without hydrogen sulphide on TSI were identified as *K. pneumoniae*.

### Genomic DNA extraction and sequencing

Isolates were stored at -80°C in skimmed milk tryptone glucose glycerol (STGG) broth until processed for DNA extraction at the University of Cape Coast Department of Biomedical Sciences’ laboratory. We used a low-cost, high-throughput method for DNA extraction from cultures grown from single colonies on MacConkey agar, as described previously [36]. DNA was quantified using the Qubit High Sensitivity DNA assay kit (Invitrogen, MA, USA). Aliquots of each DNA sample were sequenced using two approaches: (i) Oxford Nanopore MinION with R9.4.1 flow cells as described previously [36, 37], at the University of Cape Coast, Ghana; and (ii) Illumina NextSeq 500 platform (Illumina, San Diego, CA), at the Quadram Institute, UK.

### Basecalling, genome assembly and phylogenetic analysis

For the basecalling of nanopore fast5 files, we applied the ONT Guppy basecaller v4.0.14 [38] utilising the Super-accuracy model as previously described [39]. Our assembly process integrated both Nanopore and Illumina data, following the hybrid assembly protocol described in [40]. Briefly, this begins with a long-read assembly using Flye v2.9 [41], followed by long-read polishing using Medaka v1.11.1 [42, 43], and short-read polishing using Polypolish v0.5.0 [44]. We further refined the assemblies with POLCA (POLishing by Calling Alternatives) [45] a component of the MaSuRCA (Maryland Super Read Cabog Assembler) genome assembly and analysis toolkit v4.1.0 [46]. The integrity and accuracy of assemblies were validated using QUAST v5.0.0 [de6973bb] [61 (**Supplementary Figure 1**).

### Plasmid reconstruction and clustering

We employed the MOB-recon and MOB-typer modules from the MOB-suite program [47] to identify plasmids by scanning for key plasmid features such as replication genes (*rep*), mobilization proteins (relaxase), mate-pair formation (MPF), and the origin of transfer (*ori*T) among the contigs (the plasmid assemblies are available via Figshare at DOI: 10.6084/m9.figshare.24630756 and 10.6084/m9.figshare.24631020). MOB-suite assesses genomic distances against a closed plasmid reference database, and assigns plasmids to their nearest reference cluster; plasmids with genomic distances over 0.05 from the closest reference are considered novel [47]. Depending on the detected mobility genes, the suite also categorises plasmids based on their mobility potential into conjugative, mobilisable, or non-mobilisable classes.

Subsequently, we determined and visualised the common resistance genes within each cluster through a heatmap, employing the Matplotlib library in Python, indicating genes carried by plasmids in each cluster. Rare clusters with two or fewer plasmids were grouped as ‘Others’ for the purpose of visualisation.

### Genotyping and cgMLST clustering with Pathogenwatch

We uploaded our hybrid assemblies to the Pathogenwatch platform v21.3.0 [48] for comprehensive genotyping. This included *Klebsiella* species assignments, 7-gene multilocus sequence type (ST) calling [49], detection of capsular polysaccharide (K) and lipopolysaccharide (O) locus types via Kaptive v2.0.7 [50], implemented via Kleborate, and identification of acquired virulence factors and AMR determinants using Kleborate v2.3.0 [51, 52]. Kleborate assigns a virulence score ranging from 0 to 5, based on detecting specific loci (yersiniabactin, colibactin and aerobactin, in increasing order of importance) associated with increasing risk of invasiveness. Furthermore, Pathogenwatch implements the Life Identification Number (LIN) code scheme [53], facilitating the clustering of core genome MLST profiles and providing a robust method for identifying and referencing *K. pneumoniae* complex lineages.

Pathogenwatch employs a concatenated alignment of 1,972 core genes, spanning 2,172,367 bp, as a foundation for calculating pairwise single nucleotide polymorphism (SNP) distances among *K. pneumoniae* genomes. Utilising the resulting pairwise distance matrix, the platform constructs a neighbour-joining phylogenetic tree, providing insights into the genetic relationships between the bacterial isolates in a user-defined collection.

For *K. pneumoniae*, launching an isolate’s ‘Genome Report’ within the Pathogenwatch environment allows the closest relatives among all publicly available genomes to be retrieved via the “View Clusters” button, according to a user-specified threshold of alleles within the cgMLST scheme (based on a concatenated alignment of 1,972 genes (2,172,367bp) that make up the core gene library for *K. pneumoniae* in Pathogenwatch). Pathogenwatch employs a single-linkage algorithm to cluster genomes between a threshold of 0 to 50 alleles within the cgMLST scheme. Using this approach, we determined the closest neighbours to our study isolates within the context of the global *K. pneumoniae* population represented by public data available in Pathogenwatch (n=32,642 genomes as of 20^th^ December 2023).

### Data analysis and visualisation

Data analysis and visualisation involved the use of geopandas, matplotlib, and seaborn libraries in Python to map sample locations and display assembly metrics. Phylogenetic relationships were explored and visualised in R v4.1.0 using packages ggtree v3.0.4, ggplot2 v3.4.4, and phangorn v2.11.1.

### Data availability

WGS data for this study have been deposited in the National Center for Biotechnology Information (NCBI) Sequence Read Archive (SRA) under accession number PRJNA1052100 (https://shorturl.at/esWZ2). The individual accession numbers for the sequencing reads are available in **Supplementary File 3**. The interactive tree and Kleborate output are available to explore at https://microreact.org/project/7ovyxjdtatdpyfw9tdbesf.

## Ethical approval

The Institutional Review Boards of the respective laboratories granted ethical approval. Informed patient consents were waived as samples were obtained from routine diagnostics. Patient data associated with these isolates were anonymised, ensuring no possibility of patient identification based on age, sex, or hospital-related information.

## Results

### Demographic characteristics of the study population

We initially collected 159 non-duplicate isolates identified biochemically as *K. pneumoniae* from the five participating health facilities. WGS showed 45 of these isolates were non-*Klebsiella pneumoniae* (see below) or failed to meet WGS quality control standards (e.g., total genome length >7.5 Mbp or less than 4.5 Mbp), these isolates were therefore excluded from further analysis (**Figure 1B**). The final genome collection comprised isolates from 103 patients, including 49 females, 51 males, and three individuals of unspecified gender. Predominantly paediatric, 65% of the isolates were derived from patients under five years of age, 26% (n=27) from adults aged 46-65, and 9% (n=9) from over 65-year-olds. Blood and urine were the primary sources of isolates, constituting 24% (n=25) and 31% n=32), respectively (**Table 1**).

**Table 1:** Characteristics of the study population.

|  | **female**<br>**(N=49)** | **male**<br>**(N=51)** | **unknown**<br>**(N=3)** | **Total**<br>**(N=103)** |
| --- | --- | --- | --- | --- |
| **Source specimen** |  |  |  |  |
| blood | 9 (18.4%) | 16 (31.4%) | 0 (0%) | 25 (24.3%) |
| hvs | 6 (12.2%) | 0 (0%) | 0 (0%) | 6 (5.8%) |
| pus | 1 (2.0%) | 0 (0%) | 0 (0%) | 1 (1.0%) |
| sputum | 13 (26.5%) | 14 (27.5%) | 0 (0%) | 27 (26.2%) |
| swab | 3 (6.1%) | 1 (2.0%) | 0 (0%) | 4 (3.9%) |
| urine | 15 (30.6%) | 15 (29.4%) | 2 (66.7%) | 32 (31.1%) |
| wound | 2 (4.1%) | 3 (5.9%) | 0 (0%) | 5 (4.9%) |
| tracheal aspirate | 0 (0%) | 2 (3.9%) | 0 (0%) | 2 (1.9%) |
| unknown | 0 (0%) | 0 (0%) | 1 (33.3%) | 1 (1.0%) |
| **Source location** |  |  |  |  |
| Accra | 4 (8.2%) | 8 (15.7%) | 3 (33.3%) | 15 (14.6%) |
| Cape Coast | 14 (28.6%) | 17 (33.3%) | 0 (0%) | 31 (30.1%) |
| Kumasi | 12 (24.5%) | 13 (25.5%) | 0 (0%) | 25 (24.3%) |
| Sekondi-Takoradi | 19 (38.8%) | 13 (25.5%) | 0 (0%) | 32 (31.1%) |
| **Age group** |  |  |  |  |
| 0-1 | 3 (6.1%) | 6 (11.8%) | 0 (0%) | 9 (8.7%) |
| 2-5 | 28 (57.1%) | 28 (54.9%) | 2 (66.7%) | 58 (56.3%) |
| 46-55 | 9 (18.4%) | 7 (13.7%) | 0 (0%) | 16 (15.5%) |
| 56-65 | 6 (12.2%) | 4 (7.8%) | 1 (33.3%) | 11 (10.7%) |
| 66+ | 3 (6.1%) | 6 (11.8%) | 0 (0%) | 9 (8.7%) |
| **Collection year** |  |  |  |  |
| 2017 | 12 (48.0%) | 13(52.0%) | 0 (0%) | 25 (24.3%) |
| 2021 | 37 (47.4%) | 38 (48.7%) | 3 (3.8%) | 78 (75.7%) |

### Species identification and misidentification

Conventional diagnostics misidentified numerous isolates as *K. pneumoniae*. Post-sequencing verification using Pathogenwatch’s Speciator [48] revealed 103 true *K. pneumoniae,* with additional species including *K. quasipneumoniae*, *K. aerogenes*, *K. variicola*, *Escherichia coli*, *Proteus mirabilis*, *Enterobacter hormaechei*, and *E. cloacae* among others. Eleven sequences failed quality control (**Figure 1B**). The subsequent analysis focused on the confirmed *K. pneumoniae sensu stricto* isolates (n=103).

### Sequence type and K loci diversity

The n=103 *K. pneumoniae* isolates were diverse, comprising 44 unique STs (**Figure 2**) distributed across 25 different clonal groups (CGs). Remarkably, 18% (n=18) of our study isolates were assigned novel CGs, suggesting that these groups may represent emergent or region-specific lineages.

**Figure 2:**
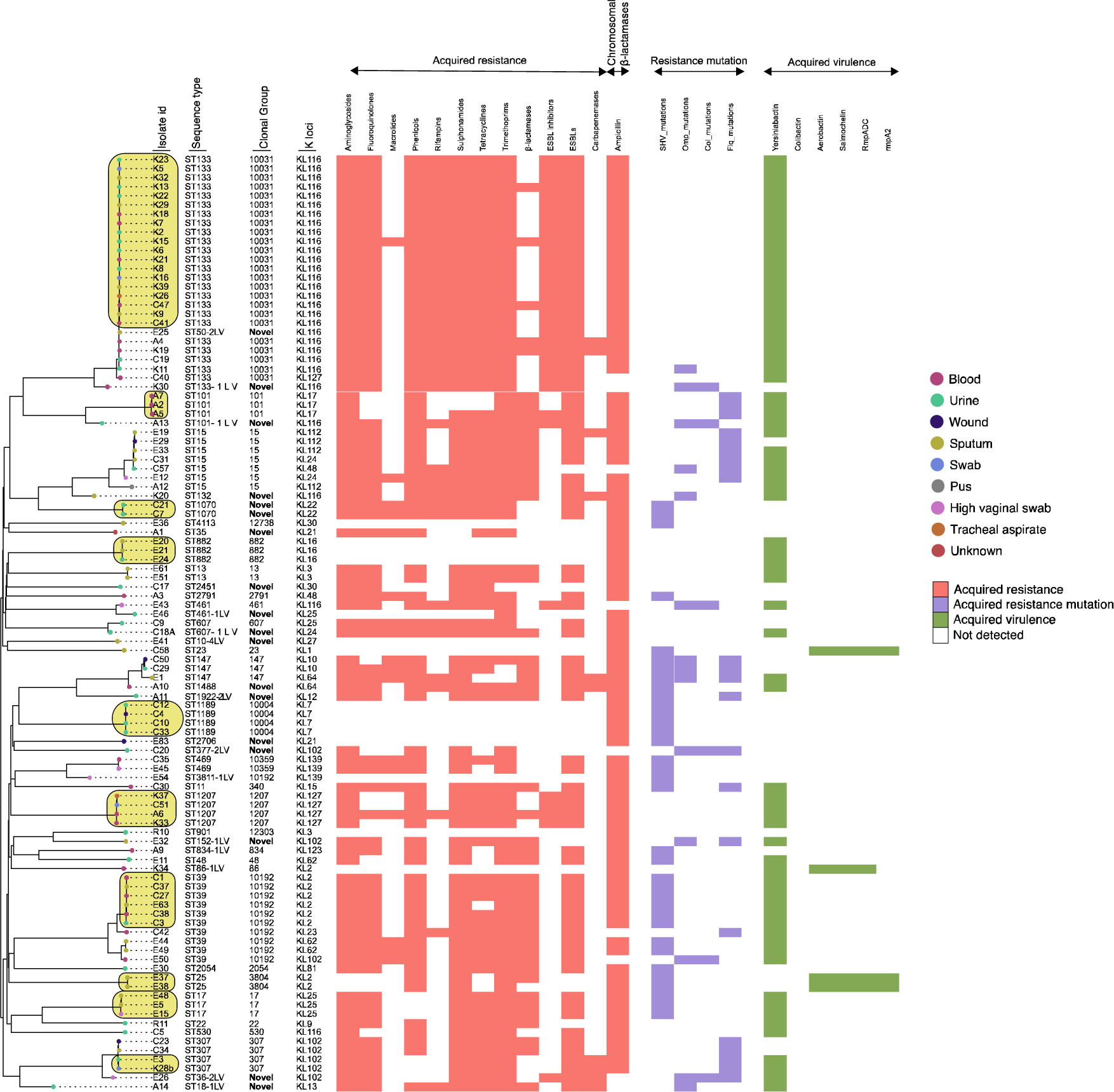
Phylogenetic analysis of the study isolates with antimicrobial resistance and virulence annotations. The figure depicts the evolutionary relationships among the study isolates, as determined by phylogenetic inference. The tree was reconstructed using the APE package via Pathogenwatch (Reference [48]), based on a concatenated alignment of 1,972 genes (2,172,367 bp) that constitute the core gene library for *K. pneumoniae* in Pathogenwatch. Each tip of the tree corresponds to a unique isolate, coloured by the source of infection (indicated in the legend), with annotations indicating the presence of acquired antimicrobial resistance genes, resistance mutations and virulence factors. Putative transmission clusters are highlighted in light khaki. The tree was rooted using the midpoint method using the phangorn package, which places the root at the midpoint of the longest distance between any two terminal nodes, balancing the tree and aiding in the interpretation of evolutionary paths. The figure was generated using the ggtree package in R and annotated using Adobe Illustrator.

ST133 (CG 10031) emerged as the predominant sequence type, accounting for n=24 (23%) of the isolates and was present in three of the four geographic regions sampled. This indicates a widespread distribution of ST133 in our study population, with potential implications for its role in disease transmission and persistence in these areas. Following closely was ST39 (CG 39), representing n=9 (9%) of the isolates and ST15 (CG 15) with n=7 (7%), while ST307 (CG 307), ST1189 (CG 10004), and ST1207 (CG 1207) each contributed n=4 isolates (4%). Geographically, Sekondi-Takoradi exhibited the highest diversity, featuring twenty-one unique STs out of n=32 isolates, followed by Cape Coast with sixteen STs out of n=31 isolates, Accra with 13 STs, and Kumasi with just six STs. Notably, no ST was detected across all four study regions, although ST1207, ST133, ST15, ST39, ST469, and ST147 were found in two or more locations. Each pair of sites shared between one and five STs (mean, 2.6).

We identified a total of 27 K locus antigens, with KL116 being the most prevalent (n=29, 28%, including n=23/29 ST133 and 6/29 other STs), followed by KL2 (n=9, 9%, including n=6/9 ST39, n=2/2 ST25 and n=1/9 ST86-1LV), KL102 (n=8, 8%, including n=4/8 ST307 and n=4 others), KL127 (n=5, all ST1207), and KL25 (n=3/3 ST17, 5% each), as well as KL112 and KL7 (4% each). These seven K loci collectively represented 64% of the study strains. Of seven O types identified, O1 (n=66/103, 64%), O2afg (n=9/103, 9%), and O2a (n=8/103, 8%) were the most frequently encountered (**Supplementary File 3)**.

### Genetic diversity of clinical K. pneumoniae from Southern Ghana

Using Pathogenwatch’s single-linkage clustering search, we identified several (n=18) isolates that were genetically distinct and lacked close relatives among publicly available genomes within the 50-allele threshold for cgMLST clustering (**Supplementary File 1**). Notably, this included isolates belonging to ST1488, ST35, ST901, ST2451, ST1070, and eleven other STs.

The dominant ST in our study, ST133 (uniformly *bla*_CTX-M-15_ positive), shared its closest genetic relation with a *bla*_CTX-M-15_ positive ESBL-producing neonatal sepsis isolate from Nigeria, as reported in the BARNARDs study [54]. This relative differed by 20 alleles within the cgMLST framework, suggesting spread within the West African region. By contrast, ST39, the second-most prevalent ST, was genetically close to isolates from Senegal and the UK, differing by merely four alleles—highlighting this clone’s regional and global dissemination. Both isolates from Senegal and the UK carried the *bla*_CTX-M-15_ gene.

Isolates of globally-disseminated multidrug-resistant clones found in our study, including ST15, ST307, ST17, ST11, ST101, and ST48, all had close relatives (≤5 allele differences) amongst public genomes from other countries and continents (**Supplementary File 1**), consistent with widespread global dissemination.

### Single nucleotide polymorphism differences in isolates within and across sampling sites

We downloaded the Pathogenwatch pairwise distance matrix, which facilitated a focused investigation into the single nucleotide polymorphisms (SNPs) present within *K. pneumoniae* genomes across the sampling sites. Our analysis revealed instances of pair-wise SNP differences less than 10 SNPs, suggestive of potential nosocomial transmission events [39, 55–57], within and between the sampling locations as follows.

Among the isolates sampled in Accra, two genomes, designated A2 and A7, displayed a pair-wise SNP difference of merely 8 SNPs, indicating a very close genetic relationship typical of recent divergence. Among the isolates from Cape Coast, a more diverse set of genomes (C10, C12, C1, C33, C4, C27, C37, and C38) showed SNP differences ranging from 4 to 9 SNPs. Similarly, in Kumasi, several genomes (K8, K9, K7, K6, K5, K39, K32, K29, K23, K21, K18, K22, K16, and K13) exhibited SNP differences between 2 and 9 SNPs, pointing to a clonal expansion likely facilitated by nosocomial vectors. A parallel trend was discerned in the following isolates from Sekondi-Takoradi, where genomes E5, E48, E15, E20, E21, E24, E37, and E38) displayed SNP differences ranging from 3 to 9 SNPs. The genetic proximity observed in these genomes exceeds the expected diversity from community-acquired strains and aligns more closely with the genomic homogeneity expected of potential nosocomial transmission clusters [39, 55–57] (**Figure 2**, highlighted).

Analysis of SNP differences between isolates from the different sites also highlighted several instances of close genetic relatedness (below the threshold of 10 SNPs), indicating potential shared or parallel sources of infection, or the movement of strains between these locations. For example, isolates E63 (from Sekondi-Takoradi) and C1 (sourced from Cape Coast) showed SNP differences as low as 3 SNPs, with similar closeness observed in pairs E63 - C27 and E63 - C38. This degree of closeness warrants further investigation into the epidemiological connections between these isolates.

Similarly, multiple isolates from Kumasi (K13, K16, K18, K21, K22, K23, K29, K32, K39, K5, K6, K7, K8 and K9) exhibited SNP differences ranging from 2 to 7, when compared to isolate C41 from Cape Coast, suggesting a cluster of closely related strains circulating within or between these sites. Additionally, isolate K37 differed from C51 by only 9 SNPs (**Figure 2**, highlighted).

### Distribution of resistance genes by antibiotic class

We identified 94 unique AMR genes spanning eleven antibiotic classes (**Supplementary File 3**). Most isolates (89/103, 86%) harboured acquired AMR genes, predominantly against aminoglycosides, trimethoprims, sulfamethoxazole (86/103, 83% each), and chloramphenicol (81/103, 79%). The ESBL gene *bla*_CTXM-15_ was present in 78% (n=80/103) of isolates (**Figure 2**, **Supplementary File 3**). Carbapenemases were identified in 10% (n=10/103) of isolates, comprising the *bla*_OXA-181_ and *bla*_NDM-1_ genes (7/10, 20% and 2/10, 80%). In all instances, the carbapenemase genes co-occurred with *bla*_CTX-M-15_. Four out of the 10 carbapenemase-positive isolates also exhibited porin mutations (including n=3/4 *bla*_OXA-181_-carrying isolates and n=1 NDM-positive isolate) and belonged to various STs, including such lineages known for multidrug resistance (MDR) as ST15, ST307, and ST147 [51, 58–60], as well as lesser-known STs like ST133, ST1488, ST18, ST36, and ST132. No convergence of acquired virulence traits associated with increased risk of invasiveness and ESBL and/or carbapenemase production was observed in our study population.

Acquired beta-lactamases such as OXA-1, CMY-2, LAP-2, TEM-1D, and SCO-1 were present in 48% (n=49), while rifamycin (*arr*-3) and macrolide resistance genes (*mph*A*, erm*B *and lsa*A), were found in 40% (n=41) and 17% (n=18) of the isolates, respectively. Tetracycline and fluoroquinolone resistance were prominent, with 76% (n=78) and 74% (n=76) of isolates harbouring resistance genes, respectively.

The bulk of the resistance genes were plasmid-borne. Of the observed resistance rates above, the following proportions were chromosomally encoded: aminoglycosides (*aac*(3)-IIa, *aac*(6’)-Ib-cr, *aad*A16, *aad*A2, *ant*(3”)-Ia, *ant*(6)-Ia, *aph*(3”)-Ib, *aph*(3’)-III, and *aph*(6)-Id), 30% (n=26/86); sulphonamides (*sul1, sul2)*, 11% (n=9/83); tetracyclines (*tet*(A), *tet*(C), *tet*(D), *tet*(G) and *tet*(M)), 13% (n=10/78); fluoroquinolones (*qnr*S1), 3% (n=2/76); macrolides (*erm*(B)), 6% (n=1/18); rifampin (*arr*-3), 2% (n=1/41); chloramphenicol (*cat*A1, *cat*A2 and *cat*B3 and *flo*R), 14% (n=11/81), trimethoprim (*dfr*A12*, dfr*A14*, dfr*A1*, dfrA8* and *dfr*A7), 7% (n=6/86) ESBL (*bla*_CTXM-15_), 10% (n=8/80), and carbapenemase (*bla*_OXA-181_), 20% (n=2/10). Intrinsic chromosomal β- lactamases (SHVs) occurred in 91% (n=94/103) of the study isolates.

Ten isolates demonstrated mutations in porin genes OmpK35 (n=8) and OmpK36 (n=2) in the absence of acquired carbapenemases. Fluoroquinolone resistance-associated mutations in the *gyr*A and *par*C genes were detected in 21% (n=22) of isolates. No determinants for colistin or tigecycline resistance were detected.

### Plasmid diversity and gene cargo

Nearly all the isolates (100/103, 97%) contained plasmids, averaging four per isolate, totalling 393 distinct plasmids harbouring 42 known replicon markers. The most common replicon markers were Col440I_1, IncFIB(K)_1_Kpn3/IncFII_1_pKP91, ColRNAI_1, IncFIA(HI1)_1_HI1/IncR_1, IncFIB(K)_1_Kpn3, and IncR_1; accounting for 50% of those detected (**Supplementary File 5**).

MOB-suite predicted 91% (n=358/393) of the identified plasmids to be mobilisable and grouped the plasmids into 118 clusters, including 16 novel clusters (**Supplementary File 6**). The distribution of these clusters varied considerably; for instance, clusters’ plasmid_AA103’ and ‘plasmid_AA274’ were each found in 36 genomes. On the other hand, several clusters (n=59), were unique to individual genomes (**Supplementary File 5**).

Median plasmid sizes ranged from 1,399 bp in ‘plasmid_AD092’ to 241,119.5 bp in ‘plasmid_AA275’, demonstrating the genetic diversity within these clusters. The ‘Others’ category, encompassing clusters with two or fewer plasmids, displayed a broad size spectrum from 1,551 bp to 252,411 bp.

We observed a heterogeneous distribution of AMR genes across plasmid clusters, with ‘plasmid_AA274’, ‘plasmid_AA277’, ‘plasmid_AA553’, and ‘plasmid_AA556’ showing a pronounced accumulation of multiple AMR genes, suggesting these may serve as AMR reservoirs. Clusters like ‘plasmid_AA406’, ‘plasmid_AE314’, and ‘plasmid_AE437’ were marked by siderophore virulence genes (**Figure 3**). Clusters’ plasmid_AA406’, ‘plasmid_AE314’ and ‘plasmid_AE437’ were characterised by the presence of siderophore virulence genes (*iro*B, *iro*C, *iro*D, *iro*N. *fyu*A, *irp*1, i*rp*2, *ybt*A, *ybt*E, *ybt*P, *yb*tQ, *ybt*T, and *ybt*U).

**Figure 3:**
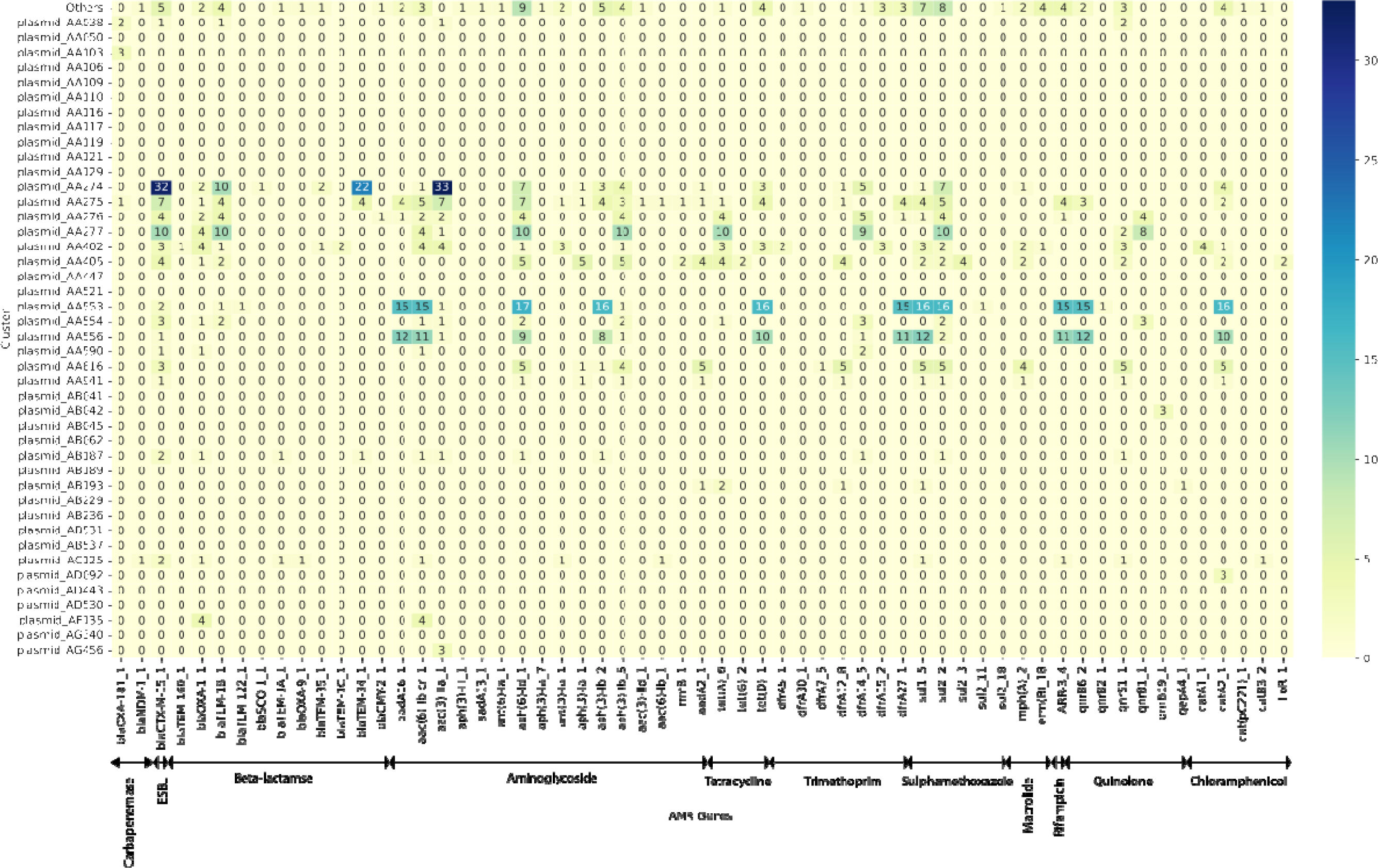
Heatmap of Antimicrobial Resistance (AMR) Genes Carried by Plasmids in Each Cluster. Each cell reflects the count of specific AMR genes found within a given cluster. The colour intensity correlates with the number of genes present, with darker shades representing higher gene counts. The scale ranges from light yellow (fewer genes) to dark blue (more genes). The numerical values in each cell denote the total count of AMR genes detected for that specific cluster. The rows are labelled with the names of plasmid clusters, while the columns correspond to specific AMR genes or groups of genes. The specific genes represented by the columns are provided in **Supplementary File 6.**

### Acquired virulence traits

The acquired virulence traits displayed less diversity compared to AMR genes. The yersiniabactin siderophore was the most frequent virulence trait, present in 70% (72/103) of the study isolates, predominantly within the *ybt*16-ICE*Kp*12 lineage (36/72, 50%). Other identified lineages identified included *ybt*10-ICE*Kp*4 (14/103, 14%), *ybt*15-ICE*Kp*11 (5/103, 5%), and *ybt*14-ICE*Kp*5 (2/102, 2%), with 6% (6/103) of isolates carrying undetermined (i.e., potentially novel) *ybt* lineages. Four isolates with hypervirulence genes were identified, these all belonged to known hypervirulent clones (n=1 ST86, n=1 ST23, n=2 ST25) (**Figure 2)**.

## Discussion

The population structure and genomic diversity of *Klebsiella pneumoniae* in Ghana, as in many sub-Saharan countries, remain poorly characterised, yet they are fundamental to the effective management and containment of infections. Our investigation sheds light on these critical aspects by analysing *K. pneumoniae* isolates from tertiary hospitals in Southern Ghana, providing new insights into their epidemiology, genetic diversity, and antimicrobial resistance. This enhanced understanding is a crucial step toward developing targeted interventions to combat the spread of this formidable pathogen.

In line with previous investigations, our study adds to the growing body of evidence indicating that conventional microbiology struggles to distinguish between members of the *K. pneumoniae* species complex [55, 61–65]. A recent study in southwestern Nigeria reported a 25% misidentification rate of *Klebsiella* as other Enterobacterales, including *Acinetobacter baumannii* and *P. aeruginosa* [66]. Our results echo these findings, underscoring the need for advanced molecular diagnostics to accurately identify pathogens and inform treatment strategies.

We have uncovered a troublingly complex picture of antimicrobial resistance in Southern Ghana, with most *K. pneumoniae* isolates harbouring multiple resistance genes. This is consistent with global trends of increasing multidrug resistance in hospital-acquired infections [1, 6, 61, 67].

The distribution of resistance genes—chromosomal versus plasmid-based—highlights the different evolutionary pressures and mechanisms at play, suggesting intrinsic resistance as well as the potential for horizontal gene transfer [68, 69].

Our data add to the mounting evidence of the *bla*_CTX-M-15_ ESBL genotype’s prevalence in healthcare and community settings in our setting [15, 16, 21, 23–25, 70–73]. The detection of carbapenem resistance genes, such as *bla*_OXA-181_ and *bla*_NDM-1_, is particularly concerning and echoes findings from other regional studies [12, 21, 24]. The presence of these genes indicates the challenging reality of treating infections with limited antimicrobial options.

The diversity of the plasmids carrying these resistance genes reflects *K. pneumoniae’s* genomic plasticity and potential to act as a reservoir for antimicrobial resistance. IncFIB and IncX3 plasmids, in particular, have been implicated in the spread of resistance, underscoring the role of mobile genetic elements in resistance gene dissemination [6, 15, 23, 24].

Our analysis also highlights the complexity of resistance mechanisms, with porin mutations contributing to carbapenem resistance [76, 77]. Although virulence traits were less diverse, the prevalence of yersiniabactin suggests it may play a significant role in the pathogenicity of *K. pneumoniae* in this region [66].

We observed a significant diversity of sequence types and K loci among *K. pneumoniae* isolates, which could inform alternative control measures like vaccines, monoclonal antibodies, and phage therapies [6, 79–82]. The predominance of specific K and O antigens may reflect their role in pathogen survival and virulence, highlighting potential targets for preventive strategies [6, 74–77].

Identifying multidrug-resistant clones such as ST15, ST307, and ST17, alongside unique genetic profiles among our isolates, points to a dynamic and evolving landscape of *K. pneumoniae* in Ghana. The disparity in sequence type distribution across various study sites reinforces the genetic diversity of these pathogens, with some sequence types being widespread while others are unique to specific locales. Previous studies corroborate our findings, with certain sequence types such as ST17 being recurrent in clinical settings and others being more geographically dispersed [18, 19, 23, 24]. This genetic variability across regions emphasises the need for localised infection control strategies, which take into account the regional differences in sequence type prevalence and the potential for localised spread of specific clones.

In a hospital setting, a genome-wide SNP difference of 21-25, corresponding to 10 Pathogenwatch SNPs, is indicative of nosocomial transmission [39, 55–57]. The observation of SNP differences less than 10 in certain pairs of genomes suggests missed opportunities in identifying and preventing hospital-acquired infections. This underscores the need for enhanced surveillance and infection control measures within hospitals and reinforce the importance of genomic surveillance in identifying potential hospital-acquired infections [66], which is paramount for informing public health interventions and antibiotic stewardship strategies.

## Limitations

This study was confined to genomic analysis, primarily due to the lack of comprehensive antimicrobial resistance (AMR) phenotype data and logistical constraints. Incorporating AMR phenotypic information, such as Minimum Inhibitory Concentrations, could have provided a richer context to the genotypic insights derived from whole-genome sequencing (WGS). Furthermore, storage complications led to the loss of some isolates, which might have resulted in an underrepresentation of the genetic diversity within the studied *K. pneumoniae* population. It is also important to note that our study encompassed only those *K. pneumoniae* isolates accessible from the participating laboratories. As some patients may have opted for private laboratory services, our findings potentially do not fully capture the breadth of *K. pneumoniae* diversity present in our study region.

## Conclusions

Our research indicates a highly diverse and antimicrobial-resistant *K. pneumoniae* population in Southern Ghana. The dominance of *bla*_CTX-M-15_ is particularly alarming, necessitating robust local and global surveillance and action. This study underscores the critical need for ongoing genomic surveillance and reinforces the importance of antimicrobial stewardship and infection prevention strategies adapted to local epidemiology.

## Acknowledgements

We extend our deepest gratitude to the laboratory personnel and administrative bodies of Korle-Bu Teaching Hospital, Greater Accra Regional Hospital (Ridge), Komfo Anokye Teaching Hospital, Cape Coast Teaching Hospital, and Effia Nkwanta Regional Hospital for their invaluable contributions to this research. Special thanks are due to Professor Zamin Iqbal and Dr Leah Roberts for their expert guidance on the plasmid cluster analysis.

## Transparency declarations

The authors declare that they have no conflicts of interest.

---

**Supplementary Figure 1:**
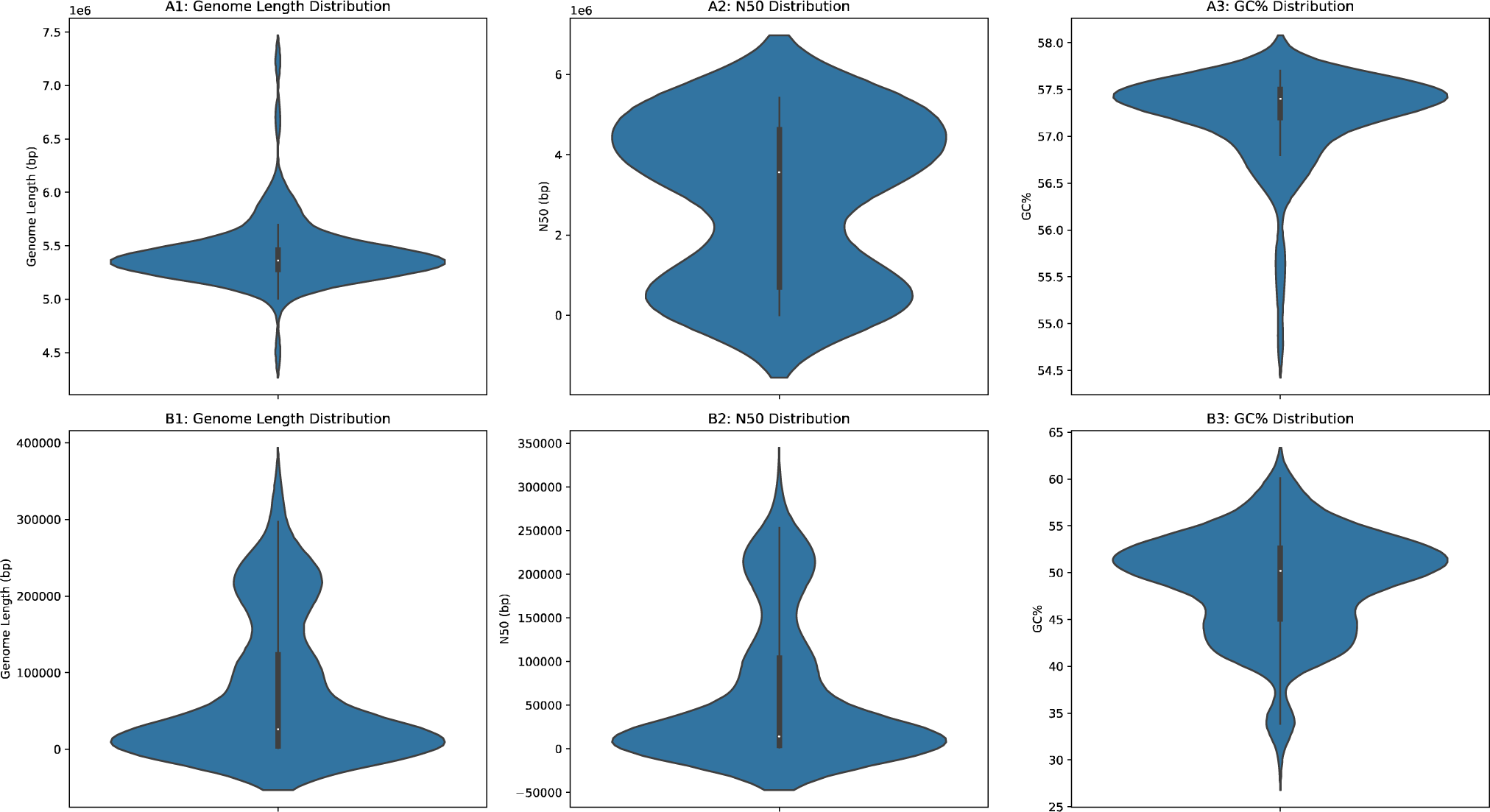
Chromosomal and plasmid assembly metrics. The figure presents the genome (**Panels A1-A3**) and plasmid (**Panels B1-B3**) assembly metrics for the *K. pneumoniae* genomes analysed in this study. The violin plots illustrate the distribution of genome lengths, N50 values, GC content percentages, and the number of contigs across multiple assemblies, highlighting the central tendency and dispersion of each metric.

